# Evaluating Plant-Microbe Associations in Response to Environmental Stressors to Enhance Salt Marsh Restoration

**DOI:** 10.1101/2024.10.14.618360

**Authors:** Kai A. Davis, Mary-Margaret McKinney, Rachel K. Gittman, Ariane L. Peralta

**Affiliations:** Department of Biology, East Carolina University, Greenville, North Carolina, USA; Integrated Coastal Sciences, East Carolina University, Greenville, North Carolina, USA; Coastal Studies Institute, Wanchese, North Carolina, USA

**Keywords:** soil microbial communities, salinity stress, microbiome stewardship, coastal protection

## Abstract

Despite substantial contributions of ecosystem services, mutualistic plant-microbe associations in wetlands are severely threatened by human activities. Therefore, promoting positive plant-microbe associations underpins coastal wetland restoration, where human stressors and climate change challenge successful outcomes. This study examined how salinity stressors influence plant-microbe relationships, where we hypothesized that the presence of microbes would provide a rescue effect by buffering abiotic stressors and yielding higher plant biomass. We used a whole sediment inocula approach and exposed marsh cordgrass (*Sporobolus alterniflorus*) plugs to a factorial experiment with three levels of microbiome addition (microbial inocula, autoclaved microbial inocula, no microbe control) and two levels of salinity (<0.5 psu, 20 psu), replicated ten times. We added marsh-site microbial inocula with autoclaved peat-based greenhouse soil and exposed half the plugs to saltwater (20 psu) and half to freshwater (<0.5 psu). Results revealed that marsh microbial inocula additions during early plant development may ameliorate salinity stressors. Plants treated with microbial inocula and salinity stress exhibited greater aboveground biomass (*P* < 0.09) than those under freshwater conditions. With live microbial inocula, the median of aboveground biomass and change in plant height were higher in saltwater compared to freshwater conditions. Change in plant height, belowground biomass, root-to-shoot ratio, soil bacterial diversity H’, and evenness J’ were similar across microbial and salinity treatments. However, salinity (*P* < 0.001) and microbial treatments (*P* < 0.001) significantly influenced the bacterial community composition, with distinct assemblages under salinity conditions. This work supports microbiome stewardship enhancing plant resiliency, improving future wetland restoration outcomes.

## INTRODUCTION

Salt marshes, coastal wetlands, and other coastal ecosystems aid in ameliorating erosion and flooding, protect people and property from storm surges, and provide habitat for fisheries and many other species (Arkema et al., 2013; Buchanan et al., 2022). Despite their high ecological and economic importance, these coastal habitats are under threat due to anthropogenic activities (e.g., land development, dredging) and climate change (e.g., accelerated sea level rise, increased mean global surface temperature, and frequency of heatwaves) (Valiela et al., 2018; Leonardi et al., 2018; Rudershausen et al., 2021; Buchanan et al., 2022; Trivedi et al., 2022). Unfortunately, current restoration efforts are failing to keep pace with current and future losses (Gittman et al. 2019), highlighting the necessity for improved salt marsh restoration techniques (Gittman et al. 2024).

The planting of marsh smooth cordgrass *Sporobolus alterniflorus* (also known as *Spartina alterniffora*) is a common approach to salt marsh restoration efforts, as the plant is a foundation species that can withstand a wide range of salinities, inundation periods, and anoxic stress (Gedan & Bertness, 2010). These foundation or habitat-forming species alter their environment, making it possible for other organisms to establish and endure within an ecosystem (Bruno, 2000). However, when commercially raised nursery plants are transplanted to field restoration sites, they are exposed to novel abiotic stressors (e.g., salinity and wave energy) and biotic factors (e.g., species interactions) that can induce stress and hinder restoration (Billah et al., 2022). Published studies evaluating the effects of exposing *S. alterniflorus* plants to abiotic stressors prior to establishment at a restoration site are lacking (but *see* Sloey et al. 2025). In addition, the understanding of plant-microbe associations between marsh soil microbiomes and *S. alterniflorus* and its influence on plant growth and survival is limited. Therefore, an examination of the effect of salinity stress and microbial inoculation on marsh plant productivity would further the understanding of ecological mechanisms and aid in restoring coastal wetland ecosystems.

In a broader context, microorganisms can enhance the nutrient acquisition of plants, support growth, and suppress diseases from pathogens (Mariotte et al., 2018; Farrer et al., 2022; Pantigoso et al., 2022; Chepsergon & Moleleki, 2023; Singh et al., 2023). In turn, plants can provide carbon resources and oxygen to root-associated microorganisms (Mariotte et al., 2018; Chepsergon & Moleleki, 2023). Human activities and climate change can and have altered these plant-microbe relationships in ways that promoted or suppressed plant growth (Cavicchioli et al., 2019; Rojas-Sánchez et al., 2025; Singh et al., 2023; Dini-Andreote, 2025). For example, anthropogenically caused eutrophication and pollution have disrupted carbon, nitrogen, and other nutrient cycles, leading to losses in microbial biodiversity and reduced fungal colonization of plants and microbial respiration (Wei et al., 2013; Cavicchioli et al., 2019; Lin et al., 2020). Global change, such as rising global surface temperatures, heatwaves, droughts, and sea level rise, can directly and indirectly modulate plant physiology, soil composition and chemistry, and microbial community assemblages, often leading to states of stress and dysbiosis (Trivedi et al., 2022; Dini-Andreote, 2025; Zabin et al., 2022). Conversely, the practice of microbiome stewardship, defined as the responsible and conscious management of microbial communities for beneficial conservation and restoration outcomes, can minimize or reverse biodiversity loss, as well as preserve key ecosystem functions and mechanisms (Peixoto et al., 2022; Choudoir et al., 2025). Growing evidence has also indicated that microbial communities can enhance plant survivorship and growth in high-stress environments, potentially aiding ecological restoration efforts (McHugh & Dighton, 2004; Mariotte et al., 2018; Li et al., 2021; Rojas-Sánchez et al., 2025; Dini-Andreote, 2025).

With the urgent need for enhanced restoration techniques, we aim to understand how to promote positive plant-microbe associations to aid in restoring coastal wetland ecosystems where human stressors and climate change challenge restoration outcomes. In this investigation, we evaluated the interactive effects of salinity stress and marsh sediment microbial inoculation on *S. alternifforus* productivity. Conducting a replicated factorial experiment with two levels of salinity and three levels of microbial inocula treatment, we measured the change in plant height and biomass (aboveground and belowground) of *S. alterniflorus* plugs after a 2-month duration. Additionally, we extracted genomic DNA from the sediment samples to characterize the soil microbiome of experimentally manipulated plants. As resident marsh microorganisms are well-adapted to a hypersaline environment, we hypothesized that *S. alterniflorus* plugs inoculated with established salt marsh sediment microbial communities would exhibit higher biomass than control plugs when subjected to salt stress.

## METHODS

### Microbiome x Salinity Stress Experimental Design

To test our hypothesis that *S. alterniflorus* plugs inoculated with restoration-area microbial communities would have higher growth rates than control plugs under salinity stress, we conducted a factorial experiment with three levels of microbiome addition (*No Addition*, *Autoclaved Inocula*, *Added Inocula*) and two levels of salinity (<0.5 psu, 20 psu), replicated ten times (*n*=60) (Figure 1). We collected marsh sediment from Hammocks Beach State Park (Swansboro, North Carolina), while the marsh plugs, *Sporobolus alterniflorus*, were purchased from a commercial plant nursery (Garner’s Landscaping, Inc., Newport, North Carolina) that routinely grows *S. alterniflorus* plugs from locally collected seeds for salt marsh restoration projects. The commercial marsh plugs were procured in 50-cell propagation trays, with one juvenile plant per 71 cm^3^ cell and filled with standard greenhouse mix (Sungro Fafard peat-based potting mix).

**Figure 1.**
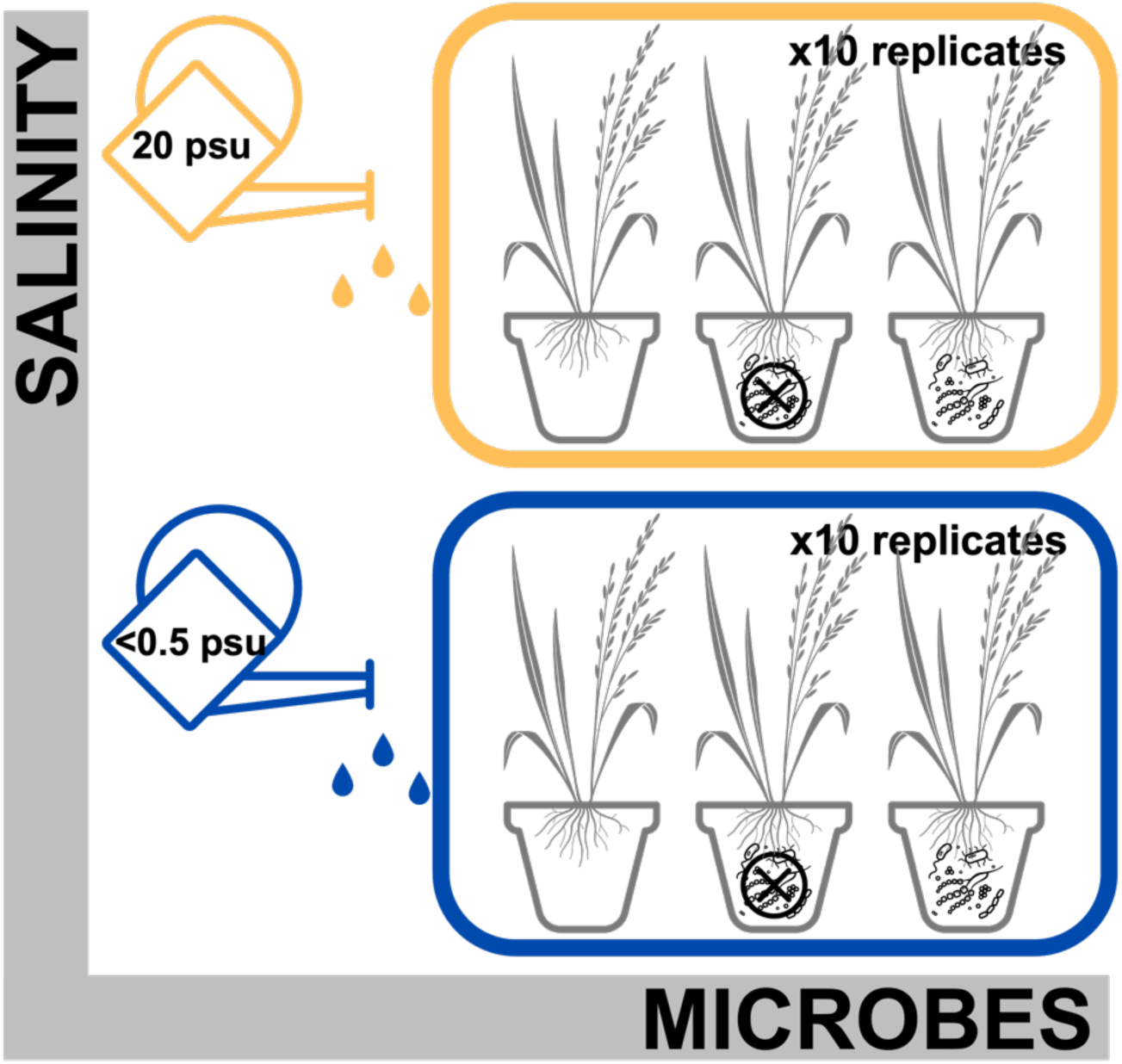
Experimental design of salinity and marsh sediment microbial inocula treatments. Factorial experiment includes three levels of microbial addition (*No Addition*, *Autoclaved Inocula*, *Added Inocula*) and two levels of salinity (<0.5 psu, 20 psu). Treatments were replicated ten times. Microbial inocula were combined with potting mix and established for four weeks, followed by weekly salt stress treatment, and harvested after eight weeks.

To prepare the microbial treatments, we combined microbial inocula with Fafard 3B potting mix, with 5% of the total volume being marsh sediment and 95% being autoclaved potting mix (following the methods of Lau & Lennon, 2011). Specifically, we combined 10 g of marsh sediment with 190 g of potting mix in an 11.25 cm square pot prior to transplanting the plant plug (25 g average weight). While autoclaving can influence the chemistry of soil organic matter and the accessibility of plant nutrients, it creates a benchmark for sterilization and microbial community establishment. Any variations observed between microbial treatments can be attributed to the introduction of inocula containing either live or dead soil microorganisms (Lau & Lennon, 2011).

The combination of untreated marsh sediment and autoclaved potting mix represented the *Added Inocula* treatment. For *Autoclaved Inocula*, marsh sediment was autoclaved before being combined, with the purpose of examining whether there were abiotic effects on plant productivity. While the microbial load may not have been completely erased, autoclaving ensured a decreased abundance of resident marsh microorganisms (31 sequence reads for autoclaved field soil compared to >26,000 sequence reads for all other sample types). For *No Addition*, 100% of the volume of the microbial treatment was comprised of autoclaved potting mix. Commercially grown *S. alterniflorus* plugs were obtained on 01 June 2023 and were transferred to 11.25 cm square pots with their respective treatments on 08 June to begin a four-week acclimation period to their new soil conditions. The plants were kept under laboratory conditions (room temperature at about 25 °C) next to a window for sunlight, and were rotated daily to ensure even light exposure.

On 23 June, the plants were transferred from the laboratory to an outside environment in Greenville, North Carolina, ∼70 miles northwest of where the plants were purchased. We placed plants on outdoor tables, in trays, where they received the same sunlight and precipitation conditions across all experimental units. We also began conducting biweekly measurements at this time, recording the maximum plant stem height (in mm) following standard methods (Currin et al., 2008; Gittman et al., 2018; Mendelssohn & Seneca, 1980; Morris & Haskin, 1990) and the number of live stems, with changes in stem height (final – initial) functioning as a proxy for growth. The four-week acclimation period concluded on 09 July, at which point the weekly salinity stress treatment was initiated and continued for four weeks.

For the salinity treatment, we bottom-watered each potted plug for 15 minutes with either ∼20 psu saltwater or ∼0.5 psu freshwater. Saltwater was made with Instant Ocean Sea Salt, with a general ratio of ∼300 g Instant Ocean to 14 L of tap water. This established a concentration of ∼20 psu, which represented salinity levels at Hammocks Beach State Park, where salinity ranges from 20 to 35 psu (personal communication, park manager). Although the duration of salt exposure in our experimental setup may be shorter compared to natural field conditions experienced by plants, it was anticipated that the deposited salt would gradually accumulate in the soil due to evaporation, mimicking real-world scenarios. This deliberate manipulation allowed us to specifically examine the effects of salt stress on plant survival and growth, distinct from the stress induced by prolonged inundation, which typically leads to anoxia. Thus, our experimental design ensured a focused investigation into salt stress dynamics without confounding stressors associated with prolonged inundation (Wang et al. 2019). Further, we opted for a short-duration, saltwater bottom-watering because it could be easily replicated by nurseries. Top-watering and subsequent freshwater flushing could be prohibitively labor- and equipment-intensive (saltwater would have to be flushed out of watering systems and equipment) as well as difficult to apply to multiple trays of plants simultaneously.

### Plant and Soil Sample Processing

At the end of the experiment, after eight weeks total, we harvested aboveground biomass and roots, dried the plant biomass samples at 60 °C for 48 hours, and weighed aboveground and root biomass. For soil bacterial characterization and analysis, we homogenized the remaining soil, transferred soil into sterile WhirlPak bags, and stored samples at -20 °C until DNA extraction and 16S rRNA amplicon sequencing (described below).

### DNA Extraction and Bacterial Amplicon Sequencing

To characterize the sediment microbiomes of the laboratory-treated plugs as well as any community composition differences, we extracted genomic DNA from sediments using the Qiagen DNeasy PowerSoil Kit, employing a combination of chemical and mechanical methods to lyse the sample following the manufacturer’s protocol. The isolated DNA was quantified using the Nanodrop spectrophotometer and diluted to a standardized concentration of 10 ng µl^-1^ to ensure uniformity across all samples. This process facilitated equimolar concentrations of DNA for downstream applications. Triplicate Polymerase Chain Reactions (PCR) were subsequently conducted on all extracted samples, serving as an efficient downstream technique for the amplified DNA. To characterize bacterial communities, we used barcoded primers (515FB/806R) initially developed by the Earth Microbiome Project (Caporaso et al. 2012) to target the V4 region of the bacterial 16S subunit of the ribosomal RNA gene (Caporaso et al., 2012; Apprill et al., 2015; Parada et al., 2016). For each sample, three 50 µL PCR libraries were prepared by combining 35.75 µL molecular grade water, 5 µL Amplitaq Gold 360 10x buffer, 5 µL MgCl2 (25 mM), 1 µL dNTPs (40mM total, 10mM each), 0.25 µL Amplitaq Gold 360 polymerase, 1 µL 515 forward barcoded primer (10 µM), 1 µL 806 reverse primer (10 µM), and 1 µL DNA template (10 ng µL^-1^). Thermocycler conditions for PCR reactions were as follows: initial denaturation (94 °C, 3 minutes); 30 cycles of 94 °C for 45 seconds, 50 °C for 30 seconds, 72 °C for 90 seconds; final elongation (72 °C, 10 minutes). The triplicate 50 µL PCR libraries were combined and then cleaned using the Axygen® AxyPrep Magnetic (MAG) Bead Purification Kit (Axygen, Union City, California, USA). Cleaned PCR products were quantified using QuantIT dsDNA BR assay (Thermo Scientific, Waltham, Massachusetts, USA) and diluted to a concentration of 10 ng µL^-1^ before pooling libraries in equimolar concentration of 5 ng µL^-1^. We sequenced the pooled libraries using the Illumina MiSeq platform using paired-end (2×250 bp) reads (Illumina Reagent Kit v2, 500 reaction kit) at the Duke University Center for Genomic and Computational Biology Core Sequencing Facility.

Sequences were processed using the mothur (v1.48.3) (Schloss et al. 2009) pipeline (Kozich et al., 2013). We assembled contigs from the paired-end reads, quality trimmed using a moving average quality score (minimum quality score 35), aligned sequences to the SILVA rRNA database (v138.2) (Quast et al. 2013), and removed chimeric sequences using the VSEARCH algorithm (Rognes et al. 2016). We created operational taxonomic units (OTUs) by first splitting sequences based on taxonomic class and then binning into OTUs based on 97% sequence similarity. Taxonomic identity was assigned using the SILVA rRNA database (v138.2) (Quast et al. 2013). We used an OTU-based 3% distance threshold instead of using amplicon sequence variant (ASV) of a single base difference, especially given our high diversity microbial communities based on short-read (250 bp) amplicon sequences, and that similar broadscale microbiome patterns are observed using OTU- or ASV-based approaches (Glassman and Martiny 2018). In addition, the OTU-based approach limits overestimation of the number of ASVs due to PCR bias and sequencing errors (Schloss 2021).

### Statistical Analyses

All statistical analyses, calculations, and graphical manipulations were run in the R environment (Rv4.5.1; R Studio 2025.05.1+513) using the vegan, ade4, picante packages, and custom functions (Dray et al., 2023; Oksanen et al., 2022). We evaluated the main and interactive effects of inoculation and salinity on *S. alterniflorus* aboveground biomass, belowground biomass, root-to-shoot ratio, and change in maximum plant stem height. We tested parametric assumptions of normality by visualizing residuals using Q-Q plots and conducting Shapiro-Wilk tests. Since the data violated the normal distribution, we conducted Kruskal-Wallis tests and *post-hoc* Dunn’s Test (Holm *P*-value adjustment, if significance was found) to assess group differences due to treatments.

Prior to microbial statistical analyses, we removed low-abundance OTUs that were represented by <10 sequences across the data set. We also omitted two samples where the total sequences were less than 10,000 reads, which represented issues with the library preparation. We calculated the relative abundance by dividing each OTU by the total number of sequence reads for each sample prior to conducting community composition analyses. Samples were organized by salinity treatment and microbial treatments. Soil bacterial diversity metrics were then computed, including Shannon Diversity (H’) and Pielou’s Evenness (J’) with corresponding ANOVA tests, providing insights into community composition by assessing both species richness and species evenness.

We evaluated the extent to which inoculation and salinity treatment explained variation in bacterial community composition using permutational multivariate analysis of variance (PERMANOVA) based on Bray-Curtis dissimilarity. We also conducted a multivariate homogeneity of group dispersions using the betadisper function in the vegan package (Table S5). We found that group dispersions according to main effects and the interaction of inoculation and salinity were similar. Therefore, significant PERMANOVA results are attributed to treatment differences and not differences in group dispersions. Next, Principal Coordinates Analysis (PCoA), based on Bray-Curtis dissimilarities, was used to visualize relationships among samples according to bacterial community composition.

To identify prominent indicator species within the bacterial communities, we used the package ‘indicspecies’. Indicator species—selected based on their ability to reflect environmental conditions—indicate changes in the environment and predict diversity patterns (De Cáceres et al., 2010). The indicspecies package facilitates this analysis by examining the relationship between species occurrence or abundance and site classification into groups representing habitat types or disturbance states (De Cáceres et al., 2010).

## RESULTS

### Marsh Microbial Addition and Salinity Stress Inffuences Plants to Varying Degrees

The effects of salinity (< 0.5 psu and 20 psu) and microbial treatments (*No Addition, Autoclaved Inocula, Added Inocula*) produced mixed results for *S. alterniflorus* growth and biomass. A notable pattern emerged where median biomass and change in plant height were higher in freshwater and saltwater for both the *No Addition* and *Autoclaved Inocula* treatments; however, this pattern was reversed for the *Added Inocula* treatment, which showed higher medians under saltwater conditions (though not always statistically significant) (Figure 2A, 2B, 2C).

**Figure 2.**
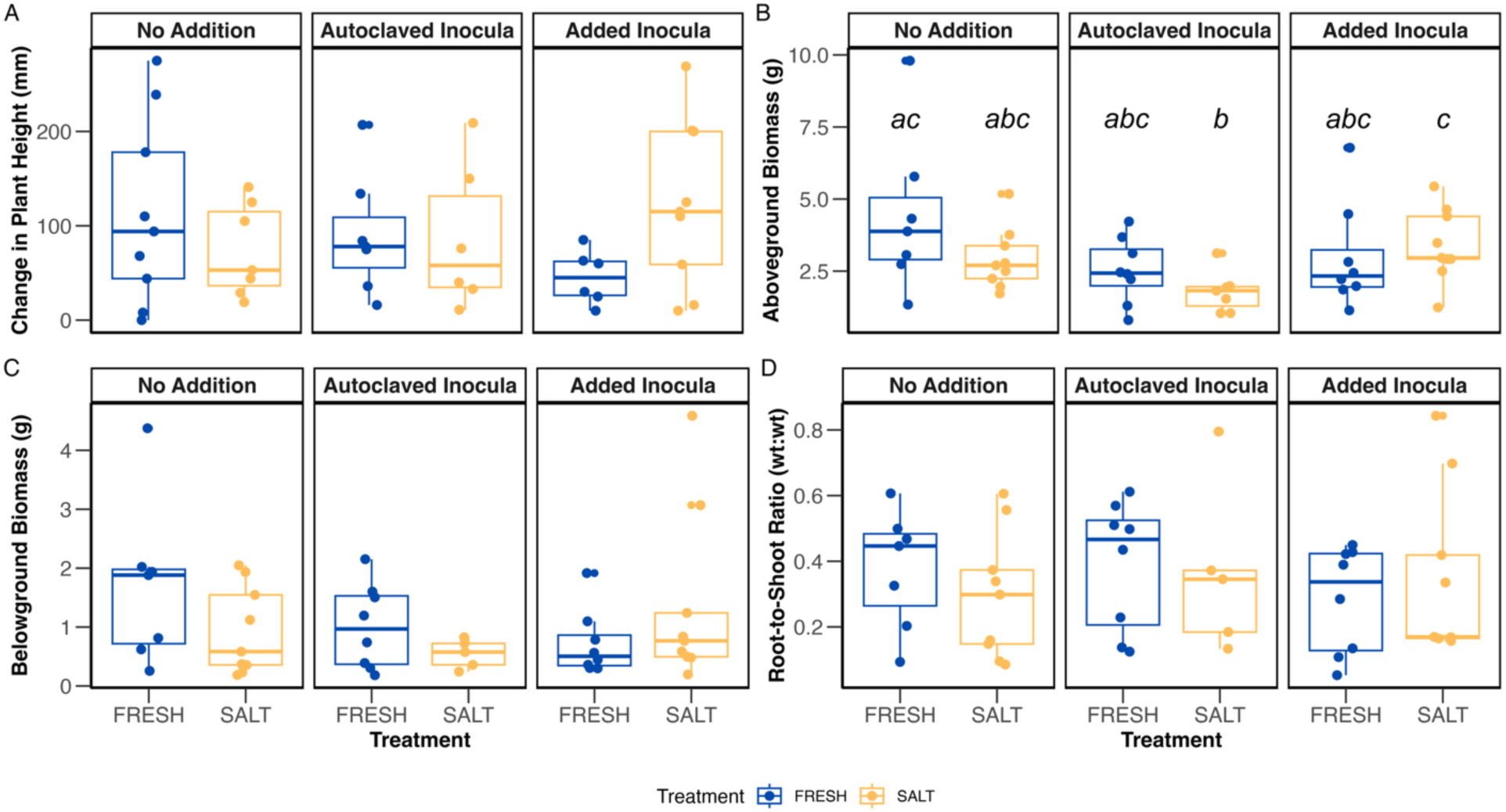
Boxplots depicting the change in plant height (in mm) **(A)**, aboveground biomass **(B)**, belowground root biomass **(C)**, and root-to-shoot ratio **(D)** on the y-axes, in response to marsh soil microbes (*No Addition, Autoclaved Inocula, Added Inocula*) and salinity stress (<0.5 psu FRESH, 20 psu SALT) treatments on the x-axes. Color represents salinity stress (orange) and freshwater control (blue), while symbols represent individual data points. The boxplot is a visual representation of 5 key summary statistics: the median, the 25% and 75% percentiles, and the whiskers, which represent the feasible range of the data as determined by 1.5 × the interquartile range. Change in plant height, belowground biomass, and root:shoot ratio results were not significant (NS). For aboveground biomass **(B)**, different letters above boxplots indicate significant difference at adjusted *P* < 0.10 based on Dunn’s test pairwise comparisons (see also Table 1).

**Table 1.** Non-parametric *post-hoc* Dunn’s test for pairwise comparisons of aboveground biomass across microbial and salinity treatments. *P*-value adjustment: Holm method. *Note*: Kruskal-Wallis test statistic: *χ* ² (5) = 11.24, *P* = 0.047.

| Comparison | Z Statistic | P-value | Adjusted P-value |
| --- | --- | --- | --- |
| No Addition Fresh – No Addition Salt | 1.18 | 0.240 | 1.00 |
| No Addition Fresh – Autoclaved Inocula Fresh | 1.68 | 0.092 | 1.00 |
| No Addition Salt – Autoclaved Inocula Fresh | 0.573 | 0.566 | 1.00 |
| No Addition Fresh – Autoclaved Inocula Salt | 2.92 | 0.003 | **0.052** |
| No Addition Salt – Autoclaved Inocula Salt | 1.92 | 0.055 | 0.709 |
| Autoclaved Inocula Fresh – Autoclaved Inocula Salt | 1.33 | 0.182 | 1.00 |
| No Addition Fresh – Added Inocula Fresh | 1.49 | 0.135 | 1.00 |
| No Addition Salt – Added Inocula Fresh | 0.372 | 0.710 | 1.00 |
| Autoclaved Inocula Fresh – Added Inocula Fresh | -0.196 | 0.844 | 0.844 |
| Autoclaved Inocula Salt– Added Inocula Fresh | -1.52 | 0.128 | 1.00 |
| No Addition Fresh – Added Inocula Salt | 0.372 | 0.710 | 1.00 |
| No Addition Salt – Added Inocula Salt | -0.859 | 0.390 | 1.00 |
| Autoclaved Inocula Fresh – Added Inocula Salt | -1.41 | 0.159 | 1.00 |
| Autoclaved Inocula Salt – Added Inocula Salt | -2.73 | 0.006 | **0.090** |
| Added Inocula Fresh – Added Inocula Salt | -1.20 | 0.228 | 1.00 |

The change in plant height (final height – initial height) over the course of the experiment was similar across salinity and microbial treatments (Kruskal-Wallis, *χ ²* (5) = 3.54, *P* = 0.617) (Figure 2A). In contrast, aboveground biomass was significantly influenced by the interaction of microbial and salinity treatments (Kruskal-Wallis, *χ ²* (5) = 11.24, *P* = 0.047). Pairwise comparisons revealed a difference in biomass between the *No Addition + Freshwater* and *Autoclaved Inocula + Saltwater* groups (Table 1, Figure 2B). The *S. alterniflorus* plants in the freshwater control had more aboveground biomass than plants in the saltwater autoclaved inocula treatment (*Z* = 2.92, *P* = 0.003, adj *P* = 0.052) (Table 1). A similar pattern was observed between *Autoclaved Inocula + Saltwater* and *Added Inocula + Saltwater* treatments, where plants with salt marsh microorganisms may have ameliorated salinity stress compared to autoclaved inocula (*Z* = -2.73, *P* = 0.006; adj *P* =0.090) (Table 1). No other pairwise comparisons were statistically significant for aboveground biomass analysis. Similarly, microbial and salinity manipulations affected belowground biomass evenly (Kruskal-Wallis, *χ* ² (5) = 4.61, *P* = 0.465) (Figure 2C). Finally, the root-to-shoot ratios were similar across treatments (Kruskal-Wallis; *χ* ² (5) = 2.65, *P* = 0.754) (Figure 2D).

### Salinity and Microbial Addition Inffuenced Bacterial Composition More Than Diversity

Salinity and microbial treatments did not influence bacterial diversity. The experimental treatments of salinity and marsh microbial inocula additions resulted in an average bacterial Shannon Diversity H’ of 6.456 ± 0.264 (Figure 3A) and Pielou’s Evenness J’ of 0.850 ± 0.018 (Figure 3B). Soil bacterial diversity metrics were similar across salinity (ANOVA, *F*1,48 = 0.035, *P* = 0.853) and microbial treatments (ANOVA, *F*2,48 = 1.53, *P* = 0.227) (Table S3). Likewise, evenness metrics were similar across treatments of salinity (ANOVA, *F*1,48 = 0.048, *P* = 0.827) and microbial inocula addition (ANOVA, *F*2,48 = 1.72, *P* = 0.190) (Table S4).

**Figure 3.**
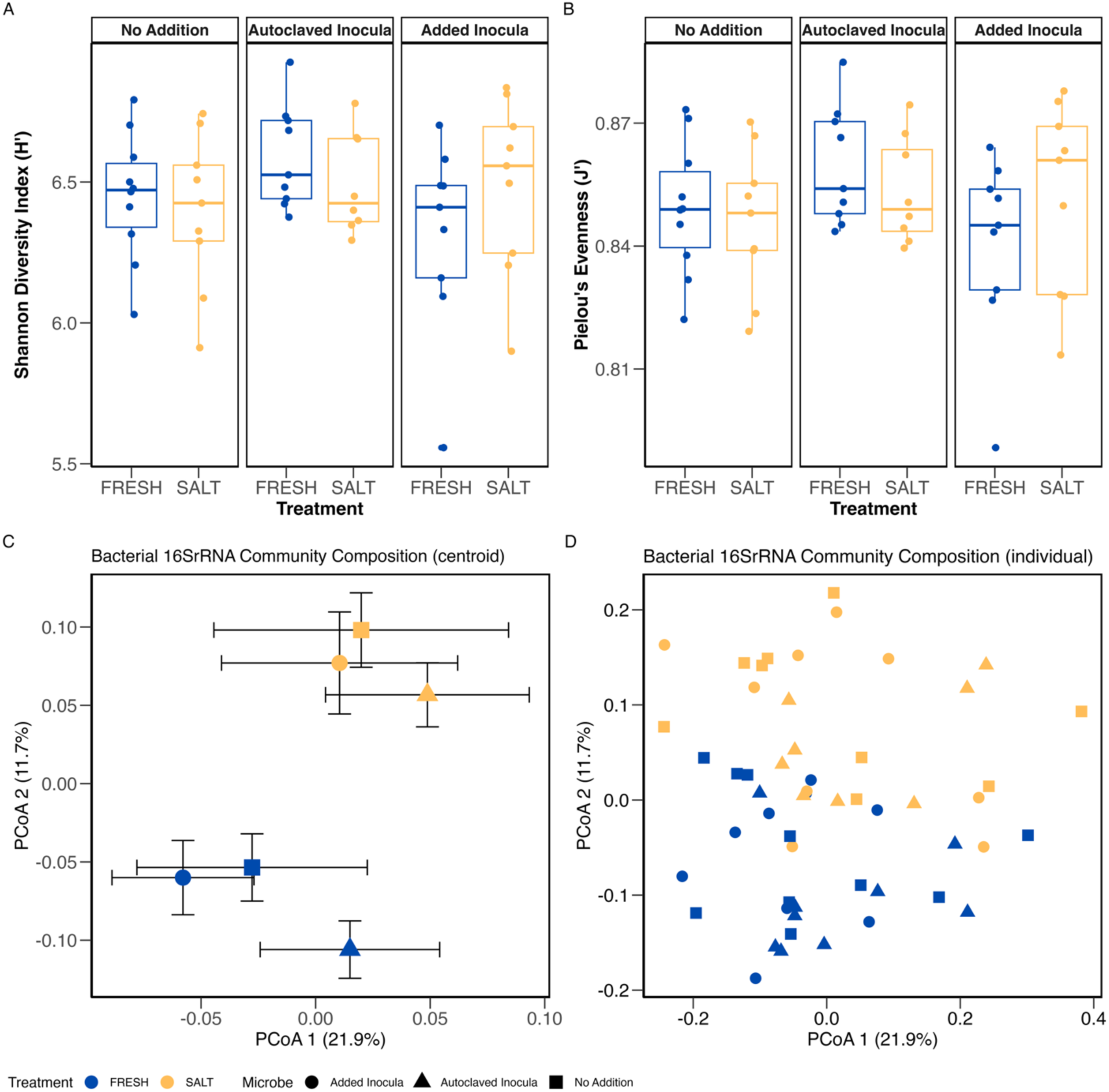
Boxplots for bacterial diversity representing Shannon Diversity Index (H’) **(A)** and Pielou’s Evenness (J’) **(B)** associated with microbial and salinity treatments. The average Shannon Diversity H’ equaled 6.456 ± 0.264, and Pielou’s Evenness J’ equaled 0.850 ± 0.018, with no significant differences found across treatments. Also included are ordination plots based on a principal coordinates analysis depicting bacterial community composition according to salinity and microbial treatments, representing the average centroid across treatments **(C)** and individual samples **(D).** Main effects of salinity (PERMANOVA: R^2^ = 0.1007, *P* < 0.001) and microbial (PERMANOVA: R^2^ = 0.0853, *P* < 0.001) treatments significantly influenced bacterial community composition. No significant effect of salinity × microbial treatment on community structure (PERMANOVA: R^2^ = 0.0396, *P* = 0.182) (See Table 2). Symbols are colored according to salinity treatment (orange = saltwater (SALT), blue = freshwater (FRESH)) and shaped by microbial treatment (circles = *Autoclaved Inocula*, triangles = *Added Inocula*, squares = *No Addition*).

**Table 2.** Summary PERMANOVA results comparing microbial community composition (based on the Bray-Curtis dissimilarity matrix) across main effects (salinity treatment, microbial inocula treatment) and the interaction of salinity × microbial treatments. Bold text indicates significant differences (*P* ≤ 0.05).

| Effect | DF | Sum of Sqs | R<sup>2</sup> | F-value | P-value |
| --- | --- | --- | --- | --- | --- |
| **Salinity TRT** | 1 | 0.4931 | 0.1007 | 6.245 | **<0.001** |
| **Microbe TRT** | 2 | 0.4176 | 0.0853 | 2.644 | **<0.001** |
| Salinity TRT × Microbe TRT | 2 | 0.1940 | 0.0396 | 1.228 | 0.182 |
| Residual | 48 | 3.7900 | 0.7743 |  |  |
| Total | 53 | 4.8946 | 1.0000 |  |  |

Conversely, microbial and salinity treatments significantly modified bacterial community composition (Table 2). The PERMANOVA, based on a Bray-Curtis dissimilarity matrix, revealed that the main effects of salinity (R^2^ = 0.1007, F = 6.245, *P* < 0.001) and microbial inocula (R^2^ = 0.0853, F = 2.644, *P* < 0.001) treatments significantly influenced bacterial community composition (Table 2). There was no significant interaction effect of salinity and microbial treatment on bacterial community structure (R^2^ = 0.0396, F = 1.228, *P* = 0.182).

To visualize the effect of treatments on bacterial community composition, we used Principal Coordinates Analysis (PCoA) based on a Bray-Curtis dissimilarity matrix (Figure 3C, 3D). We observed distinct bacterial community composition according to saltwater or freshwater exposure along the second principal coordinate axis (PCoA axis 2, accounting for 11.7% of variance). A more nuanced separation by microbial inocula was observed along the first principal coordinate axis (PCoA axis 1, accounting for 21.9% of variance). Within both salinity groups, communities showed a consistent gradient along PCoA axis 1. *Autoclaved Inocula* treatment resulted in the most distinct communities, particularly under freshwater conditions, where it was most separated from the other groups.

### Context-Dependent Responses of Bacterial Indicator Taxa Across Treatments

To further examine bacterial associations with microbial and salinity treatments, we employed indicator species analysis to identify a subset of bacterial taxa that represent the interaction and main effects of microbial and salinity treatments (Table S1). There were 46 OTUs significantly associated with freshwater conditions across all inocula treatments from phyla *Proteobacteria*, *Acidobacteria, Actinobacteria*, *Verrucomicrobia*, *Planctomycetes*, and *Gemmatimonadetes*; and 37 OTUs were associated with all saltwater treatments from phyla *Proteobacteria*, *Bacteroidetes*, *Acidobacteria*, *Verrucomicrobia,* and *Spirochaetes*. In freshwater treatments, prominent OTUs represented orders *Myxococcales*, *Caulobacterales*, *Rhizobiales,* and *Rhodospirillales*, and the genus *Opitutus*. In contrast, in saltwater treatments, top OTUs represented taxa from the orders *Myxococcales, Burkholderiales,* and *Sphingobacteriales*, family *Desulfuromonadaceae*, and genus *Dongia* (Table S1).

For the microbial treatments, 28 OTUs were significantly associated with the *No Addition* control treatment from phyla *Proteobacteria, Planctomycetes*, *Bacteroidetes*, *Acidobacteria*, *Verrucomicrobia*, *and Gemmatimonadetes*; 8 OTUs were associated with *Autoclaved Inocula* from phyla *Proteobacteria* and *Verrucomicrobia*; and 24 OTUs were associated with *Added Inocula* from phyla *Proteobacteria*, *Verrucomicrobia*, *Acidobacteria,* and *Actinobacteria* (Table S2). The top OTUs for the *No Addition* treatments represented taxa from the order *Myxococcales*, families *Plantomycetaceae*, *Chitnophagaceae,* and genera *Rhizomicrobium* and *Opitutus*. For *Autoclaved Inocula*, indicator species represented bacterial taxa in the orders *Myxococcales* and *Rhodospirillales*, taxa in the family *Comamonadaceae,* and taxa in the genus *Hyphomicrobium*. Finally, top indicator species for *Added Inocula* represented taxa in the genera *Geobacter*, *Prosthecobacter, Phenylobacterium,* and *Opitutus,* and taxa in the class *Gp1* (Acidobacteria) (Table S2).

## DISCUSSION

While understanding the underpinnings of microbial ecology is still in its relative infancy compared to more contemporary scientific disciplines, the roles of belowground microbial communities in coastal wetland restoration have been of great interest (Birnbaum & Trevathan-Tackett, 2023). Many empirical studies have demonstrated the reliance of plants on soil microbiomes in tolerating environmental stresses, including salinity, inundation, drought, and limited nutrient availability (reviewed in Farrer et al., 2022; Rodriguez et al., 2008). Furthermore, the adaptive and evolutionary responses of plants to changing environmental conditions have been shown to rely heavily on the context of microbial community assemblage, with plants exhibiting increased fitness when associated with stress-adapted microbiomes (Lau & Lennon, 2012). Presently, gaps in knowledge hinder the success of complete ecosystem restoration projects (Duarte et al., 2015); therefore, the foundational knowledge gained from biological conservation and species restoration studies is of utmost relevance and importance. In our investigation, the interactive effects of microbial inocula additions and salinity treatment on *S. alterniflorus* height, biomass, and bulk soil microbiomes produced various results. We hypothesized that inoculating *S. alternifforus* plugs with salt marsh microbial communities would buffer salt stress and promote plant growth; a subset of our results supports this hypothesis.

The effect of salinity on plant growth depended on the microbial treatment. The addition of live marsh microbes resulted in higher median plant biomass and changes in plant height under saltwater compared to freshwater, while the opposite pattern was observed for plants receiving autoclaved microbes or the no addition control. Despite these observable trends, statistical analysis concluded that there was no significant effect of microbial and salinity treatments on the change in plant height and belowground biomass. This null result, particularly for bioinoculation experiments, is not uncommon across both laboratory and field settings and can be attributed to several factors (reviewed in Compant et al., 2019). Possible explanations include, but are not limited to, insufficient establishment of microbial communities due to the short 8-week duration of our design; absent vital satellite taxa (i.e., rare taxa) in the given dosage of microbial inocula; and/or the multifactorial nature of salinity stress limiting the rescue effect of inocula (e.g., alleviating osmotic stress, but not ion toxicity), resulting in a plant growth response too subtle to detect statistically without more precise physiological metrics and/or larger sample size. Recent evidence has also indicated that endophytic bacteria (those within the plant tissue) provide greater benefits to host health when under stress than rhizome-associated bacteria (Malik & Arora, 2022; Peng et al., 2023). In all, the high variability observed in plant growth responses within treatment groups likely overwhelmed the signal of any potential treatment effect, reducing the statistical power of our experiment.

The findings from this study obscure a potentially important biological pattern within our figures. Specifically, *Added Inocula* was the only microbial treatment group where saltwater displayed a greater median of change in height than those in freshwater (Figure 2A). While this pairwise comparison (*Added Inocula + Salt* versus *Added Inocula + Fresh*) was not significant (Dunn’s Test; *P* = 0.07 before adjustment), the reversal of the predicted negative effects of salinity stress on plant growth is noteworthy and may hint at amelioration. To further investigate whether marsh microbial inoculation ameliorates abiotic stress, we recommend future experiments with greater statistical power (increased sample sizes) to address variability in responses, as well as more enhanced physiological stress markers (e.g., proline accumulation, leaf ion concentration responses to stress) (Hayat et al., 2012; Azizi et al., 2017).

Aboveground biomass results provided more nuance and clarity on the interaction of salinity and microbial additions to plant growth. Compared to freshwater control, *Autoclaved Inocula* under salinity stress experienced a greatly inhibited biomass (Table 1, Figure 2B). These results provide evidence that the sterilization of belowground microbial communities and the addition of chemically inert soil components (i.e., sand) can hinder plant productivity, possibly inducing nutrient limitations for the host plants and nutrient cycling microbes. Comparatively, plants under salinity stress that were inoculated with marsh microorganisms exhibited greater aboveground biomass, further supporting a growing body of research that microbial inocula additions during early plant development may be critical for establishing resilience to environmental stressors (Trivedi et al., 2021; Peng et al., 2023; Shade, 2023) For example, greenhouse-raised *S. alterniflorus* that were exposed to saltwater (10 ppt salinity) and microbial consortium (a selection of plant growth-promoting rhizobacteria) experienced the highest rates of growth when compared to other treatments (Bledsoe & Boopathy, 2016).

Belowground biomass and root-to-shoot ratios were similar across treatment types (Figure 2C, 2D), providing further insight into the plants’ stress response strategy and possible limitations of our methodology. The short duration of our experiment may have been insufficient to elicit measurable changes in biomass allocation. Longer-term morphological changes, like root system restructuring, are often not as prioritized as immediate physiological adjustments (i.e., shoot growth) (Poorter et al., 2012). Additionally, the physical constraints of the pot sizes may have limited root expansion and shoot growth, possibly masking treatment effects. Despite this, some studies have suggested that optimal growth levels can be reached over a broad range of carbon allocation patterns (Sugiura & Tateno, 2011). As *S. alterniflorus* plants under *Added Inocula + Salt* exhibited the lowest root:shoot median compared to other treatments (Figure 2D), a compelling hypothesis for our results is that the beneficial microorganisms (i.e., plant-growth-promoting rhizobacteria) did not change the plant’s allocation of carbon but rather improved the efficiency of its use (Yang et al., 2009). Future work should investigate how pre-exposing *S. alterniflorus* to salinity stress and marsh microbiomes influences long-term productivity after transplantation to field conditions. In a natural, unrestricted setting, the evaluation of carbon allocation between roots and shoots would be more ecologically relevant than in the restrictive environment of a pot.

Salinity stress and microbial inocula additions reshaped bacterial community composition without altering overall diversity or evenness. When analyzing patterns in bacterial diversity, average alpha diversity metrics Shannon H’ and Pielou’s evenness J’ were similar across microbial and salinity treatments (Table S3 & S4). *Autoclaved Inocula* treatment tended to decrease the ranges of Shannon diversity (H’) and evenness (J’), while greater ranges existed in the *Added Inocula* (Figure 3A, 3B). Additionally, marsh microbial inocula under salinity stress tended to exhibit greater medians in Shannon diversity (H’) and evenness (J’) than those of freshwater, in support of the reversal trend. Past studies have shown that increases in microbial biodiversity and evenness are associated with increased resistance to plant pathogens, improved soil health and ecosystem functions, increased plant growth and survivorship, and many other positive benefits (see Banerjee & van der Heijden, 2023; Birnbaum & Trevathan-Tackett, 2023; Farrer et al., 2022; Graham & Knelman, 2023; Lau & Lennon, 2012). Despite similar bacterial diversity, microbial and salinity treatments significantly influenced community composition, with distinct bacterial communities forming in the salt and freshwater groups (Table 2, Figure 3C, 3D). This may indicate the strength of the selective filtering of environmental conditions in microbial community assembly, rather than acting as forces that uniformly suppressed or enhanced microbial richness and evenness. As the composition of microorganisms mattered more than their sheer diversity in the context of specific environmental filters, this should be highly considered when applying microbial inocula for restoration.

As microbial sampling techniques and high-throughput sequencing continue to improve, increasing evidence has highlighted the utility of microbial communities as indicator species (Liang et al., 2023; Ma et al., 2022; Urakawa & Bernhard, 2017). Microorganisms can function as rapid and sensitive bioindicators and exhibit patterns of spatial distribution (e.g., distinct rhizosphere and bulk soil communities) (Urakawa & Bernhard, 2017; Wang et al., 2016); our findings contribute to this growing body of literature. While indicator species analysis revealed dominant microbial assemblages from phyla *Proteobacteria*, *Acidobacteria*, *Actinobacteria*, *Bacteroidetes*, and *Verrucomicrobia*—phyla known to be relatively abundant and ubiquitous in soil (Janssen, 2006)— there were clear distinctions in OTUs with little overlap between treatments (Table S1 & S2). Seven OTUs from the family *Opitutaceae* (phylum *Verrucomicrobia*) overlapped in *Added Inocula*, saltwater, and freshwater treatments, which have been found in a wide variety of environments but have been associated with nutrient cycling in low-oxygen areas (Jayakumar et al., 2017). At the phylum/class level, many indicator OTUs from *Added Inocula* and saltwater were classified into *Verrucomicrobia, Acidobacteria GP1,* and *Alphaproteobacteria*, all of which are acknowledged to contain well-known halotolerant and/or oligotrophic bacteria (Kielak et al., 2016; Zhang et al., 2024). Out of ten OTUs classified into *Bacteroidetes*, 70% were found exclusively in saltwater treatments, supporting evidence of halo-tolerance (Canfora et al., 2014), while interestingly, the remaining three were found in the *No Addition* group. In general, however, microbial bioindicators were unique among salinity and microbial addition treatments, further providing evidence that plant-microbe interactions and environmental conditions uniquely structure soil communities (Li et al., 2021; Zhang et al., 2024). It is important to note that the majority of environmental microbes remain unculturable and unidentified (Stewart, 2012), which represent 20 bacterial taxa that could not be resolved past the phylum or kingdom levels. Moreover, microorganisms in nature often live in complex, synergistic communities, with the vast majority of functions and interactions remaining out of current technological and scientific grasp (Widder et al., 2016).

The patterns in bacterial community composition revealed support for strong environmental filtering due to salinity exposure, even under short (15-minute) weekly pulses. Bacterial community composition was relatively similar among microbial treatments but revealed distinct bacterial assemblages when under the influence of salinity treatments. As such, the pre-treatment of saline conditions and resident microbial inocula may provide rescue effects from abiotic and biotic field stressors after marsh grass plugs are transplanted and exposed to ex situ environmental conditions. This work provides evidence that microbial stewardship is essential for buffering against ecological stressors and could promote successful plant establishment for wetland restoration. Additionally, our work supports the theory of “habitat-adapted symbioses,” highlighting the facultative mutualisms between marsh plants and their symbiotic microorganisms in tolerating high-stress environments (Rodriguez et al., 2008). With many stressors projected to intensify in the near future (e.g., extreme climatic events, sea level rise, droughts and heatwaves, compounding stress from invasive species, anthropogenic change, etc.), marshes will need to adapt to survive, and their associated microorganisms will undergo even more rapid evolutionary changes (Zabin et al., 2022; Billah et al., 2022; Lau & Lennon, 2012; Mueller et al., 2020). By incorporating microbial inocula into salt marsh restoration practices, success could be better leveraged.

Within a broader scope, coastal wetlands and salt marshes are among the most productive ecosystems on Earth and provide numerous ecological and economic services. Salt marshes support fisheries and habitats for biodiversity, filter contaminants, sequester carbon, buffer storm surges and protect shorelines from erosion, provide tourism, and facilitate many other functions (Arkema et al., 2013; Buchanan et al., 2022). The preservation and restoration of these habitats are thus vital, aligning with the principles of the One Health framework, which emphasizes how the health of humans, wildlife, and the environment is inextricably linked (Trinh et al., 2018; Peixoto et al., 2022; Banerjee & van der Heijden, 2023). By evaluating the interactions between *Sporobolus alternifforus*, marsh sediment microbiomes, and environmental stressors, this study provides insight into ecological dynamics and may aid in salt marsh restoration efforts.

## Acknowledgments

We acknowledge the staff at Hammocks Beach State Park and Garner’s Landscaping for their assistance in conducting the field experiment. We thank Gittman lab members Jennifer Fickler and James Kelley for assistance in setting up and maintaining the field research, and Peralta lab members Toby Gallegos, Daphka Joseph, Scott Siebor, Dejah Smith, and Colin Finlay for lab assistance. We also thank Aeran Coughlin and Jean-Philippe Gibert for microbiome sequencing support. This work was supported by the National Science Foundation (#1845845, #2009185, # 2302609 to A.L.P.), the U.S. Coastal Research Program and the National Oceanographic and Atmospheric Administration National Sea Grant (#2023-0534-02/22-NC-USCRP-3 to R.K.G.) and North Carolina Sea Grant (#PAM-P24-002985-SA-02 to R.K.G. and A.L.P.) and East Carolina University (Undergraduate Research and Creative Activity Award to K.A.D.).

## STATEMENTS AND DECLARATIONS

The authors declare no competing interests.

