## Supplemental Information (v3) for "Evaluating Plant-Microbe Associations in Response to Environmental Stressors to Enhance Salt Marsh Restoration"

**RUNNING TITLE: Salt marsh plant-microbiome restoration**

Kai A. Davis[1], Mary-Margaret McKinney[2], Rachel K. Gittman[1], Ariane L. Peralta[1,*]

[1]Department of Biology, [2]Integrated Coastal Sciences, East Carolina University, Greenville, North Carolina, USA, 3 Coastal Studies Institute, Wanchese, North Carolina, USA

**Table S1.** Bacterial taxa (OTUs) representing the unique taxa associated with the freshwater or saltwater treatment according to indicator species analysis. This summary represents the top bacterial taxa associated with each treatment type. Blank spaces represent unclassified taxonomy.

| OTU | Cluster | IndVal | Prob | Domain | Phylum | Class | Order | Family | Genus |
|---|---|---|---|---|---|---|---|---|---|
| Otu00215 | Freshwater | 0.822 | 0.001 | Bacteria | Proteobacteria | | | | |
| Otu00051 | Freshwater | 0.783 | 0.001 | Bacteria | Verrucomicrobia | Opitutae | Opitutales | Opitutaceae | Opitutus |
| Otu00049 | Freshwater | 0.766 | 0.001 | Bacteria | Verrucomicrobia | Opitutae | Opitutales | Opitutaceae | Opitutus |
| Otu00203 | Freshwater | 0.742 | 0.001 | Bacteria | Verrucomicrobia | Subdivision3 | | | |
| Otu00031 | Freshwater | 0.736 | 0.001 | Bacteria | Proteobacteria | Deltaproteobacteria | Myxococcales | Polyangiaceae | |
| Otu00275 | Freshwater | 0.734 | 0.001 | Bacteria | | | | | |
| Otu00255 | Freshwater | 0.708 | 0.011 | Bacteria | Proteobacteria | Gammaproteobacteria | | | |
| Otu00131 | Freshwater | 0.696 | 0.001 | Bacteria | Proteobacteria | Betaproteobacteria | | | |
| Otu00246 | Freshwater | 0.692 | 0.003 | Bacteria | Proteobacteria | Alphaproteobacteria | Rhodospirillales | | |
| Otu00195 | Freshwater | 0.689 | 0.001 | Bacteria | | | | | |
| Otu00102 | Freshwater | 0.665 | 0.001 | Bacteria | Proteobacteria | Alphaproteobacteria | Caulobacterales | Caulobacteraceae | Phenylobacterium |
| Otu00067 | Freshwater | 0.659 | 0.003 | Bacteria | Proteobacteria | Deltaproteobacteria | | | |
| Otu00291 | Freshwater | 0.653 | 0.001 | Bacteria | Proteobacteria | Alphaproteobacteria | Caulobacterales | Caulobacteraceae | Phenylobacterium |
| Otu00108 | Freshwater | 0.652 | 0.001 | Bacteria | Proteobacteria | Alphaproteobacteria | Rhizobiales | | |
| Otu00356 | Freshwater | 0.649 | 0.015 | Bacteria | Planctomycetes | Planctomycetacia | Planctomycetales | Planctomycetaceae | Gemmata |
| Otu00285 | Freshwater | 0.643 | 0.002 | Bacteria | Proteobacteria | Alphaproteobacteria | Caulobacterales | Caulobacteraceae | |
| Otu00205 | Freshwater | 0.643 | 0.001 | Bacteria | Proteobacteria | Alphaproteobacteria | Caulobacterales | Caulobacteraceae | Phenylobacterium |
| Otu00089 | Freshwater | 0.642 | 0.001 | Bacteria | Acidobacteria | Gp6 | | | |
| Otu00158 | Freshwater | 0.635 | 0.001 | Bacteria | Proteobacteria | Alphaproteobacteria | Rhizobiales | | |
| Otu00164 | Freshwater | 0.634 | 0.029 | Bacteria | Actinobacteria | Actinobacteria | Actinomycetales | Thermomonosporaceae | Actinomadura |

| | | | | | | | | | |
|---|---|---|---|---|---|---|---|---|---|
| Otu00020 | Freshwater | 0.631 | 0.001 | Bacteria | Acidobacteria | Gp6 | | | |
| Otu00064 | Freshwater | 0.629 | 0.001 | Bacteria | Proteobacteria | Alphaproteobacteria | | | |
| Otu00070 | Freshwater | 0.627 | 0.001 | Bacteria | Proteobacteria | Alphaproteobacteria | | | |
| Otu00083 | Freshwater | 0.627 | 0.001 | Bacteria | | | | | |
| Otu00126 | Freshwater | 0.627 | 0.004 | Bacteria | | | | | |
| Otu00094 | Freshwater | 0.625 | 0.002 | Bacteria | Proteobacteria | Alphaproteobacteria | Rhodospirillales | | |
| Otu00165 | Freshwater | 0.621 | 0.001 | Bacteria | Proteobacteria | Alphaproteobacteria | Rhizobiales | | |
| Otu00202 | Freshwater | 0.618 | 0.004 | Bacteria | Proteobacteria | Alphaproteobacteria | Rhodospirillales | | |
| Otu00052 | Freshwater | 0.617 | 0.011 | Bacteria | Actinobacteria | Actinobacteria | Solirubrobacterales | | |
| Otu00138 | Freshwater | 0.617 | 0.015 | Bacteria | Proteobacteria | Deltaproteobacteria | Myxococcales | | |
| Otu00075 | Freshwater | 0.614 | 0.001 | Bacteria | Proteobacteria | Betaproteobacteria | | | |
| Otu00039 | Freshwater | 0.610 | 0.002 | Bacteria | Proteobacteria | Alphaproteobacteria | | | |
| Otu00022 | Freshwater | 0.609 | 0.001 | Bacteria | Actinobacteria | Actinobacteria | Solirubrobacterales | | |
| Otu00218 | Freshwater | 0.603 | 0.001 | Bacteria | Gemmatimonadetes | Gemmatimonadetes | Gemmatimonadales | Gemmatimonadaceae | Gemmatimonas |
| Otu00002 | Freshwater | 0.600 | 0.001 | Bacteria | Proteobacteria | Alphaproteobacteria | ***Incertae sedis*** | ***Incertae sedis*** | Rhizomicrobium |
| Otu00235 | Freshwater | 0.597 | 0.022 | Bacteria | Verrucomicrobia | Verrucomicrobiae | Verrucomicrobiales | Verrucomicrobiaceae | Prosthecobacter |
| Otu00035 | Freshwater | 0.594 | 0.002 | Bacteria | Proteobacteria | Alphaproteobacteria | Rhizobiales | Bradyrhizobiaceae | Bradyrhizobium |
| Otu00032 | Freshwater | 0.592 | 0.002 | Bacteria | Proteobacteria | Alphaproteobacteria | Rhizobiales | | |
| Otu00298 | Freshwater | 0.591 | 0.009 | Bacteria | Proteobacteria | | | | |
| Otu00019 | Freshwater | 0.585 | 0.003 | Bacteria | Proteobacteria | Betaproteobacteria | | | |
| Otu00197 | Freshwater | 0.580 | 0.005 | Bacteria | Proteobacteria | Alphaproteobacteria | | | |
| Otu00178 | Freshwater | 0.580 | 0.024 | Bacteria | Actinobacteria | Actinobacteria | Actinomycetales | Micromonosporaceae | Rugosimonospora |
| Otu00087 | Freshwater | 0.579 | 0.012 | Bacteria | Actinobacteria | Actinobacteria | Actinomycetales | Acidothermaceae | Acidothermus |
| Otu00013 | Freshwater | 0.560 | 0.013 | Bacteria | Proteobacteria | Alphaproteobacteria | Rhizobiales | | |
| Otu00023 | Freshwater | 0.553 | 0.031 | Bacteria | Proteobacteria | Alphaproteobacteria | Rhodospirillales | | |
| Otu00011 | Freshwater | 0.548 | 0.003 | Bacteria | Proteobacteria | Alphaproteobacteria | Rhizobiales | Hyphomicrobiaceae | Devosia |
| Otu00191 | Saltwater | 0.912 | 0.001 | Bacteria | Proteobacteria | Deltaproteobacteria | Myxococcales | | |
| Otu00176 | Saltwater | 0.812 | 0.001 | Bacteria | Bacteroidetes | | | | |

| | | | | | | | | | |
|---|---|---|---|---|---|---|---|---|---|
| Otu00170 | Saltwater | 0.810 | 0.001 | Bacteria | Proteobacteria | Betaproteobacteria | Burkholderiales | | |
| Otu00133 | Saltwater | 0.773 | 0.001 | Bacteria | | | | | |
| Otu00154 | Saltwater | 0.754 | 0.001 | Bacteria | Proteobacteria | Alphaproteobacteria | Rhodospirillales | Rhodospirillaceae | Dongia |
| Otu00040 | Saltwater | 0.726 | 0.005 | Bacteria | Proteobacteria | Deltaproteobacteria | Desulfuromonadales | Desulfuromonadaceae | |
| Otu00147 | Saltwater | 0.718 | 0.003 | Bacteria | Proteobacteria | Deltaproteobacteria | Myxococcales | | |
| Otu00110 | Saltwater | 0.703 | 0.001 | Bacteria | Acidobacteria | Gp1 | | | |
| Otu00062 | Saltwater | 0.688 | 0.001 | Bacteria | Bacteroidetes | Sphingobacteria | Sphingobacteriales | Chitinophagaceae | |
| Otu00128 | Saltwater | 0.681 | 0.001 | Bacteria | Acidobacteria | Gp1 | | | |
| Otu00260 | Saltwater | 0.677 | 0.004 | Bacteria | Proteobacteria | | | | |
| Otu00219 | Saltwater | 0.666 | 0.006 | Bacteria | Proteobacteria | Deltaproteobacteria | Myxococcales | | |
| Otu00090 | Saltwater | 0.661 | 0.05 | Bacteria | Bacteroidetes | Sphingobacteria | Sphingobacteriales | Rhodothermaceae | Salinibacter |
| Otu00092 | Saltwater | 0.652 | 0.001 | Bacteria | Verrucomicrobia | Subdivision3 | | | |
| Otu00093 | Saltwater | 0.652 | 0.002 | Bacteria | Verrucomicrobia | Subdivision3 | | | |
| Otu00236 | Saltwater | 0.643 | 0.008 | Bacteria | | | | | |
| Otu00111 | Saltwater | 0.638 | 0.007 | Bacteria | Proteobacteria | Betaproteobacteria | Burkholderiales | Comamonadaceae | Hydrogenophaga |
| Otu00184 | Saltwater | 0.637 | 0.036 | Bacteria | Bacteroidetes | Sphingobacteria | | | |
| Otu00163 | Saltwater | 0.633 | 0.008 | Bacteria | Spirochaetes | Spirochaetes | Spirochaetales | Spirochaetaceae | Spirochaeta |
| Otu00283 | Saltwater | 0.629 | 0.018 | Bacteria | Bacteroidetes | Sphingobacteria | Sphingobacteriales | Chitinophagaceae | |
| Otu00190 | Saltwater | 0.628 | 0.001 | Bacteria | Verrucomicrobia | Opitutae | Opitutales | Opitutaceae | Opitutus |
| Otu00063 | Saltwater | 0.624 | 0.005 | Bacteria | Verrucomicrobia | Subdivision3 | | | |
| Otu00230 | Saltwater | 0.622 | 0.002 | Bacteria | Verrucomicrobia | Opitutae | Opitutales | Opitutaceae | Alterococcus |
| Otu00257 | Saltwater | 0.616 | 0.007 | Bacteria | Acidobacteria | Gp3 | | | |
| Otu00279 | Saltwater | 0.611 | 0.004 | Bacteria | Proteobacteria | Deltaproteobacteria | Myxococcales | | |
| Otu00081 | Saltwater | 0.609 | 0.043 | Bacteria | Bacteroidetes | Sphingobacteria | Sphingobacteriales | Chitinophagaceae | |
| Otu00233 | Saltwater | 0.607 | 0.011 | Bacteria | Proteobacteria | Betaproteobacteria | | | |
| Otu00144 | Saltwater | 0.602 | 0.039 | Bacteria | Proteobacteria | Alphaproteobacteria | Rhodospirillales | | |
| Otu00206 | Saltwater | 0.601 | 0.01 | Bacteria | Proteobacteria | Alphaproteobacteria | Rhodospirillales | Rhodospirillaceae | Dongia |
| Otu00001 | Saltwater | 0.595 | 0.002 | Bacteria | Bacteroidetes | Sphingobacteria | Sphingobacteriales | Chitinophagaceae | |

| | | | | | | | | | |
|---|---|---|---|---|---|---|---|---|---|
| Otu00045 | Saltwater | 0.590 | 0.017 | Bacteria | Proteobacteria | Gammaproteobacteria | Xanthomonadales | Sinobacteraceae | Steroidobacter |
| Otu00232 | Saltwater | 0.583 | 0.039 | Bacteria | Acidobacteria | Gp3 | | | |
| Otu00014 | Saltwater | 0.573 | 0.016 | Bacteria | | | | | |
| Otu00033 | Saltwater | 0.570 | 0.045 | Bacteria | Proteobacteria | Gammaproteobacteria | | | |
| Otu00159 | Saltwater | 0.569 | 0.025 | Bacteria | Proteobacteria | Deltaproteobacteria | | | |
| Otu00025 | Saltwater | 0.559 | 0.044 | Bacteria | Proteobacteria | Alphaproteobacteria | Rhodospirillales | Rhodospirillaceae | Dongia |
| Otu00018 | Saltwater | 0.558 | 0.005 | Bacteria | Acidobacteria | Gp6 | | | |

**Table S2.** Bacterial taxa (OTUs) representing the unique taxa associated with the microbial addition treatment according to indicator species analysis. This summary represents the top bacterial taxa associated with each treatment type. Blank spaces represent unclassified taxonomy.

| OTU | Cluster | IndVal | Prob | Domain | Phylum | Class | Order | Family | Genus |
|---|---|---|---|---|---|---|---|---|---|
| Otu00215 | No Addition | 0.707 | 0.001 | Bacteria | Proteobacteria | | | | |
| Otu00356 | No Addition | 0.609 | 0.001 | Bacteria | Planctomycetes | Planctomycetacia | Planctomycetales | Planctomycetaceae | Gemmata |
| Otu00240 | No Addition | 0.529 | 0.002 | Bacteria | Proteobacteria | Alphaproteobacteria | ***Incertae sedis*** | ***Incertae sedis*** | Rhizomicrobium |
| Otu00283 | No Addition | 0.493 | 0.006 | Bacteria | Bacteroidetes | Sphingobacteria | Sphingobacteriales | Chitinophagaceae | |
| Otu00061 | No Addition | 0.483 | 0.001 | Bacteria | Proteobacteria | Alphaproteobacteria | | | |
| Otu00314 | No Addition | 0.482 | 0.018 | Bacteria | Proteobacteria | Deltaproteobacteria | Myxococcales | | |
| Otu00076 | No Addition | 0.455 | 0.001 | Bacteria | Acidobacteria | Gp3 | | | |
| Otu00322 | No Addition | 0.453 | 0.069 | Bacteria | Verrucomicrobia | Opitutae | Opitutales | Opitutaceae | Opitutus |
| Otu00063 | No Addition | 0.447 | 0.028 | Bacteria | Verrucomicrobia | Subdivision3 | | | |
| Otu00104 | No Addition | 0.445 | 0.001 | Bacteria | Proteobacteria | Alphaproteobacteria | | | |
| Otu00081 | No Addition | 0.445 | 0.079 | Bacteria | Bacteroidetes | Sphingobacteria | Sphingobacteriales | Chitinophagaceae | |
| Otu00186 | No Addition | 0.445 | 0.001 | Bacteria | Proteobacteria | Alphaproteobacteria | Rhizobiales | | |
| Otu00101 | No Addition | 0.442 | 0.001 | Bacteria | Acidobacteria | Gp6 | | | |
| Otu00106 | No Addition | 0.442 | 0.002 | Bacteria | Proteobacteria | Alphaproteobacteria | Rhodospirillales | | |
| Otu00251 | No Addition | 0.442 | 0.001 | Bacteria | Proteobacteria | Alphaproteobacteria | Rhodospirillales | Rhodospirillaceae | |
| Otu00067 | No Addition | 0.440 | 0.055 | Bacteria | Proteobacteria | Deltaproteobacteria | | | |
| Otu00039 | No Addition | 0.437 | 0.002 | Bacteria | Proteobacteria | Alphaproteobacteria | | | |
| Otu00212 | No Addition | 0.408 | 0.073 | Bacteria | Acidobacteria | Gp1 | | | |
| Otu00014 | No Addition | 0.406 | 0.019 | Bacteria | | | | | |

| | | | | | | | | | |
|---|---|---|---|---|---|---|---|---|---|
| Otu00291 | No Addition | 0.400 | 0.055 | Bacteria | Proteobacteria | Alphaproteobacteria | Caulobacterales | Caulobacteraceae | Phenylobacterium |
| Otu00002 | No Addition | 0.400 | 0.025 | Bacteria | Proteobacteria | Alphaproteobacteria | ***Incertae sedis*** | ***Incertae sedis*** | Rhizomicrobium |
| Otu00210 | No Addition | 0.399 | 0.097 | Bacteria | Gemmatimonadetes | Gemmatimonadetes | Gemmatimonadales | Gemmatimonadaceae | Gemmatimonas |
| Otu00025 | No Addition | 0.398 | 0.024 | Bacteria | Proteobacteria | Alphaproteobacteria | Rhodospirillales | Rhodospirillaceae | Dongia |
| Otu00001 | No Addition | 0.396 | 0.058 | Bacteria | Bacteroidetes | Sphingobacteria | Sphingobacteriales | Chitinophagaceae | |
| Otu00311 | No Addition | 0.395 | 0.1 | Bacteria | | | | | |
| Otu00197 | No Addition | 0.387 | 0.073 | Bacteria | Proteobacteria | Alphaproteobacteria | | | |
| Otu00082 | No Addition | 0.378 | 0.029 | Bacteria | Proteobacteria | Gammaproteobacteria | Xanthomonadales | Xanthomonadaceae | Dokdonella |
| Otu00010 | No Addition | 0.364 | 0.074 | Bacteria | Proteobacteria | Alphaproteobacteria | | | |
| Otu00255 | Autoclaved Inocula | 0.527 | 0.096 | Bacteria | Proteobacteria | Gammaproteobacteria | | | |
| Otu00155 | Autoclaved Inocula | 0.447 | 0.028 | Bacteria | Proteobacteria | Betaproteobacteria | | | |
| Otu00048 | Autoclaved Inocula | 0.443 | 0.017 | Bacteria | Verrucomicrobia | | | | |
| Otu00138 | Autoclaved Inocula | 0.438 | 0.032 | Bacteria | Proteobacteria | Deltaproteobacteria | Myxococcales | | |
| Otu00144 | Autoclaved Inocula | 0.426 | 0.075 | Bacteria | Proteobacteria | Alphaproteobacteria | Rhodospirillales | | |
| Otu00115 | Autoclaved Inocula | 0.419 | 0.002 | Bacteria | Proteobacteria | Betaproteobacteria | Burkholderiales | Comamonadaceae | |
| Otu00126 | Autoclaved Inocula | 0.412 | 0.078 | Bacteria | | | | | |
| Otu00073 | Autoclaved Inocula | 0.408 | 0.089 | Bacteria | Proteobacteria | Alphaproteobacteria | Rhizobiales | Hyphomicrobiaceae | Hyphomicrobium |
| Otu00012 | Added Inocula | 0.529 | 0.007 | Bacteria | Proteobacteria | Deltaproteobacteria | Desulfuromonadales | Geobacteraceae | Geobacter |
| Otu00235 | Added Inocula | 0.511 | 0.001 | Bacteria | Verrucomicrobia | Verrucomicrobiae | Verrucomicrobiales | Verrucomicrobiaceae | Prosthecobacter |
| Otu00024 | Added Inocula | 0.493 | 0.001 | Bacteria | Acidobacteria | Gp1 | | | |
| Otu00041 | Added Inocula | 0.486 | 0.001 | Bacteria | Acidobacteria | Gp1 | | | |
| Otu00226 | Added Inocula | 0.479 | 0.003 | Bacteria | Proteobacteria | Alphaproteobacteria | Caulobacterales | Caulobacteraceae | Phenylobacterium |
| Otu00092 | Added Inocula | 0.470 | 0.004 | Bacteria | Verrucomicrobia | Subdivision3 | | | |
| Otu00172 | Added Inocula | 0.469 | 0.037 | Bacteria | | | | | |
| Otu00044 | Added Inocula | 0.469 | 0.014 | Bacteria | Verrucomicrobia | Opitutae | Opitutales | Opitutaceae | Opitutus |
| Otu00105 | Added Inocula | 0.458 | 0.002 | Bacteria | Proteobacteria | Alphaproteobacteria | Sphingomonadales | Erythrobacteraceae | |
| Otu00128 | Added Inocula | 0.450 | 0.042 | Bacteria | Acidobacteria | Gp1 | | | |
| Otu00222 | Added Inocula | 0.443 | 0.044 | Bacteria | Actinobacteria | Actinobacteria | | | |

| | | | | | | | | | |
|---|---|---|---|---|---|---|---|---|---|
| Otu00110 | Added Inocula | 0.440 | 0.069 | Bacteria | Acidobacteria | Gp1 | | | |
| Otu00195 | Added Inocula | 0.430 | 0.071 | Bacteria | | | | | |
| Otu00072 | Added Inocula | 0.425 | 0.019 | Bacteria | Proteobacteria | Betaproteobacteria | Burkholderiales | Comamonadaceae | |
| Otu00285 | Added Inocula | 0.421 | 0.05 | Bacteria | Proteobacteria | Alphaproteobacteria | Caulobacterales | Caulobacteraceae | |
| Otu00043 | Added Inocula | 0.419 | 0.013 | Bacteria | Proteobacteria | Gammaproteobacteria | | | |
| Otu00232 | Added Inocula | 0.418 | 0.058 | Bacteria | Acidobacteria | Gp3 | | | |
| Otu00098 | Added Inocula | 0.418 | 0.07 | Bacteria | Verrucomicrobia | Opitutae | Opitutales | Opitutaceae | Opitutus |
| Otu00054 | Added Inocula | 0.414 | 0.038 | Bacteria | Proteobacteria | | | | |
| Otu00339 | Added Inocula | 0.412 | 0.054 | Bacteria | Verrucomicrobia | Subdivision3 | | | |
| Otu00066 | Added Inocula | 0.400 | 0.034 | Bacteria | Proteobacteria | | | | |
| Otu00135 | Added Inocula | 0.398 | 0.079 | Bacteria | Verrucomicrobia | Subdivision3 | | | |
| Otu00139 | Added Inocula | 0.397 | 0.019 | Bacteria | Proteobacteria | Alphaproteobacteria | ***Incertae sedis*** | ***Incertae sedis*** | Rhizomicrobium |
| Otu00102 | Added Inocula | 0.391 | 0.095 | Bacteria | Proteobacteria | Alphaproteobacteria | Caulobacterales | Caulobacteraceae | Phenylobacterium |

**Table S3.** Summary ANOVA comparing changes in bacterial Shannon Diversity H' due to main effects (salinity treatment, microbial inocula treatment) and the interaction salinity × microbial treatments. Bold text indicates significant differences ($P \leq 0.05$).

| **Effect** | ***DF*** | **Sum of Sqs** | **Mean Sq** | ***F-value*** | ***P-value*** |
|---|---|---|---|---|---|
| Salinity TRT | 1 | 0.002 | 0.00237 | 0.035 | 0.853 |
| Microbe TRT | 2 | 0.209 | 0.10437 | 1.530 | 0.227 |
| Salinity TRT × Microbe TRT | 2 | 0.168 | 0.08423 | 1.234 | 0.300 |
| Residual | 48 | 3.275 | 0.06824 | | |

**Table S4.** Summary ANOVA comparing changes in bacterial Pielou's Evenness J' due to main effects (salinity treatment, microbial inocula treatment) and the interaction salinity × microbial treatments. Bold text indicates significant differences (*P* ≤ 0.05).

| **Effect** | ***DF*** | **Sum of Sqs** | **Mean Sq** | ***F-value*** | ***P-value*** |
|---|---|---|---|---|---|
| Salinity TRT | 1 | 0.000016 | 0.0000161 | 0.048 | 0.827 |
| Microbe TRT | 2 | 0.001148 | 0.0005738 | 1.721 | 0.190 |
| Salinity TRT × Microbe TRT | 2 | 0.000630 | 0.0003152 | 0.945 | 0.396 |
| Residual | 48 | 0.016002 | 0.0003334 | | |

**Table S5.** Adonis2 and betadisper results for community composition and dispersion tests. Bold text indicates significant differences (*P* ≤ 0.05).

| **Factor** | **$R^2$** | ***F-value*** | ***P-value*** | **Test** |
|---|---|---|---|---|
| **Salinity TRT** | 0.101 | 6.24 | **0.001** | adonis2 |
| **Microbe TRT** | 0.085 | 2.64 | **0.001** | adonis2 |
| Salinity TRT: Microbe TRT | 0.040 | 1.23 | 0.168 | adonis2 |
| Salinity TRT | | 1.05 | 0.344 | betadisper |
| Microbe TRT | | 0.71 | 0.483 | betadisper |
| Salinity TRT: Microbe TRT | | 0.56 | 0.731 | betadisper |
